# *Liguleless narrow* and *narrow odd dwarf* act in overlapping pathways to regulate maize development & physiology

**DOI:** 10.1101/2022.03.20.485070

**Authors:** María Jazmín Abraham-Juárez, Michael Busche, Alyssa A. Anderson, China Lunde, Jeremy Winders, Shawn A. Christensen, Charles T. Hunter, Sarah Hake, Jacob O. Brunkard

**Affiliations:** Laboratorio Nacional de Genómica para la Biodiversidad, Unidad de Genómica Avanzada, Centro de Investigación y de Estudios Avanzados del Instituto Politécnico Nacional, Guanajuato 36821, México; Laboratory of Genetics, University of Wisconsin, Madison, WI 53706, USA; Department of Plant and Microbial Biology, University of California, Berkeley, CA 94720, USA; Plant Gene Expression Center, USDA Agricultural Research Service, Albany, CA 94710, USA; Genomics and Bioinformatics Research Unit, U.S. Department of Agriculture-Agricultural Research Service, Raleigh, NC, USA; Nutrition, Dietetics, and Food Science, Brigham Young University, Provo, UT 84602, USA; Chemistry Research Unit, USDA Agricultural Research Service, Gainesville, FL 32608, USA

## Abstract

*narrow odd dwarf* (*nod*) and *Liguleless narrow* (*Lgn*) are pleiotropic maize mutants that both encode plasma membrane proteins, cause similar developmental patterning defects, and constitutively induce stress signaling pathways. To investigate how these mutants coordinate maize development and physiology, we screened for protein interactors of NOD by affinity purification. LGN was identified by this screen as a strong candidate interactor, and we confirmed the NOD-LGN molecular interaction through orthogonal experiments. We further demonstrated that LGN, a receptor-like kinase, can phosphorylate NOD *in vitro*, hinting that they could act in intersecting signal transduction pathways. To test this hypothesis, we generated *Lgn-R;nod* mutants in two backgrounds (B73 and A619) and found that these mutations enhance each other, causing more severe developmental defects than either single mutation on its own, with phenotypes including very narrow leaves, increased tillering, and failure of the main shoot. Transcriptomic and metabolomic analyses of the single and double mutants in the two genetic backgrounds revealed widespread induction of pathogen defense genes and a shift in resource allocation away from primary metabolism in favor of specialized metabolism. These effects were similar in each single mutant and heightened in the double mutant, leading us to conclude that NOD and LGN act cumulatively in overlapping signaling pathways to coordinate growth-defense tradeoffs in maize.

## INTRODUCTION

Plants regularly develop new organs from meristems through ordered processes of cell expansion and division (Sluis and Hake, 2015; Richardson *et al*., 2021). Because development continues throughout the plant lifespan, plants must coordinate growth and developmental patterning with dynamic changes in their environment and physiology. This coordination is often conceptualized as “growth-defense tradeoffs” (Huot *et al*., 2014; Guo, Major and Howe, 2018; Monson *et al*., 2022), and can involve metabolic reorganization to emphasize either primary metabolism (when conditions are favorable) or secondary/specialized metabolism to produce defense-related compounds (when experiencing stress). For example, in response to pathogen attack, plants often temporarily arrest new growth in favor of synthesizing compounds that deter the pathogen (especially various phytoalexins) or even trigger localized programmed cell death to limit the infection (Bomblies and Weigel, 2007; Szczesny *et al*., 2010; Ahuja, Kissen and Bones, 2012; Li *et al*., 2012; Karasov *et al*., 2017). This hypothesis that growth and defense are intertwined is supported by discoveries at the cellular and molecular levels demonstrating that the signal transduction pathways coordinating development and stress responses often overlap (Wu *et al*., 2020). For instance, plant genomes encode hundreds of cell surface receptor-like proteins (RLPs) and receptor-like kinases (RLKs) that can be stimulated by pathogen-associated molecules, hormones, or secreted peptides, which then act through similar or convergent downstream kinase signaling cascades (especially MAP kinase transduction networks) to modulate nuclear gene expression (Perraki *et al*., 2018; Albert *et al*., 2020; Dievart *et al*., 2020; Gou and Li, 2020; DeFalco and Zipfel, 2021). As another example, calcium (Ca^2+^) is trafficked across the membranes by diverse transporters, including the mechanosensitive PIEZO proteins, nucleotide-binding leucine rich-repeat receptors (NLRs), and cyclic nucleotide-gated channels (CNGC), which variously promote growth, activate defense responses while inhibiting growth, or trigger programmed cell death, despite all fundamentally acting as Ca^2+^ transporters (Bi *et al*., 2021; Jacob *et al*., 2021; Mousavi *et al*., 2021; Radin *et al*., 2021; Xu *et al*., 2022). Deciphering how these complex signal transduction pathways intersect and diverge is a major goal for plant biology, especially in agricultural crops, where breeding robust and resilient crop plants that can withstand stresses without sacrificing yields will benefit from a detailed mechanistic understanding of kinase, calcium, and other signaling networks (Liu *et al*., 2017; Bailey-Serres *et al*., 2019; Brunkard, 2020; Karavolias *et al*., 2021; Luan and Wang, 2021; Lutt and Brunkard, 2022).

From forward genetic screens for maize mutants defective in ligule patterning, we characterized two mutants, *Liguleless narrow (Lgn)* and *narrow odd dwarf (nod)*, that act at the intersection of plant development and stress responses (Moon, Candela and Hake, 2013; Rosa *et al*., 2017; Anderson *et al*., 2019). NOD encodes a putative mechanosensitive Ca^2+^-permeable channel (Nakagawa *et al*., 2007; Furuichi *et al*., 2012; Rosa *et al*., 2017; Yoshimura, Iida and Iida, 2021). The Arabidopsis orthologues of NOD, MID1-COMPLEMENTING ACTIVITY 1 (MCA1) and MCA2, are proposed to participate in various responses to environmental stimuli, but *mca1;mca2* mutants do not show severe development defects under standard growing conditions (Yamanaka *et al*., 2010; Mori *et al*., 2018). In contrast, *nod* mutants are defective in organogenesis and cellular differentiation, with phenotypic severity depending on inbred background (Rosa *et al*., 2017). In B73, the boundary between blade and sheath is obliterated, leaves are narrow, and cell division and differentiation are abnormal in *nod* mutants. These plants are also highly tillered, and their primary shoots fail to develop. In A619, mutants appear similar to wild-type as seedlings, but at maturity, *nod* mutants are shorter with narrower leaves. In Mo17, *nod* mutants are nearly indistinguishable from wild-type, showing only a modest decrease in height (Rosa *et al*., 2017).

LGN encodes a plasma membrane-localized receptor-like kinase (Moon, Candela and Hake, 2013). The reference allele, *Lgn-R*, carries a semidominant missense mutation that abolishes its kinase activity. Like *nod*, the *Lgn-R* mutation causes pleiotropic phenotypes that express differently depending on its genetic background (Buescher *et al*., 2014; Anderson *et al*., 2019). In B73, heterozygotes have narrow leaves, lack a ligule at the leaf margins, often lack ears, and have fewer branches in the tassel (Moon, Candela and Hake, 2013). *Lgn-R* mutants in B73 are also sensitive to temperature, with higher temperatures often causing lethality (Anderson *et al*., 2019). *Lgn-R/Lgn-R* homozygotes resemble *nod* mutants in B73. In A619 and Mo17, *Lgn-R* heterozygotes have a normal ligule and near-normal height. We used the difference in phenotype between Mo17 and B73 in a QTL mapping experiment to identify a genetic modifier of *Lgn-R*, which we named *Sympathy for the ligule (SOL)* (Buescher *et al*., 2014). *SOL e*ncodes an orthologue of the Arabidopsis MAP3K-inhibiting protein, ENHANCED DISEASE RESISTANCE 4 (EDR4) (Anderson *et al*., 2019). Phosphoproteomic and transcriptomic analyses suggested that disruption of LGN activity in *Lgn-R* triggers a MAP kinase signaling cascade that constitutively activates biotic stress responses in these mutants, but the *SOL* allele in the Mo17 background can rescue this phenotype, perhaps by driving sufficient *SOL* expression to repress the hyperactivated MAP kinase signaling cascade.

Here, we sought to understand how NOD impacts development and immunity in plants by discovering its potential protein interactors. To our surprise, we identified LGN as a potential NOD-interacting protein. Using a range of biochemical, molecular, cellular, genetic, genomic, and metabolomic approaches, we interrogated the relationship between NOD and LGN, demonstrating that they act through overlapping pathways to impact maize physiology and development.

## RESULTS

### Protein-protein interaction screens identified LGN as an interactor of NOD

To begin to unravel how NOD impacts maize development and stress responses, we took a molecular approach by screening for potential protein interactors of NOD that could participate in NOD-dependent signaling pathways. We took two orthogonal approaches to screen for NOD interactors: co-immunoprecipitation (coIP) and yeast two-hybrid (Y2H) screens. Using a polyclonal antibody that specifically recognizes NOD (Rosa *et al*., 2017), NOD was immunoprecipitated from three-week old wild-type B73 seedlings (Fig. 1A). *nod* B73 mutants lacking NOD protein were used as a negative control to account for any nonspecific immunoprecipitates. As expected, NOD protein was purified and strongly enriched in eluates after immunoprecipitation of wild-type B73 protein extracts, but NOD was not present in *nod* mutant eluates (Fig. 1A). Tubulin was detected in initial protein extracts and in flow-through, but not in the final wash nor in the eluate, demonstrating that abundant, nonspecific interactors were effectively removed during the co-immunoprecipitation protocol. After these quality control steps, immunoprecipitates were analyzed by mass spectrometry to identify potential NOD interacting partners. This experiment was repeated three times, with slight modification in one replicate to enrich for membrane proteins by adding phosphatase inhibitors and increasing the concentration of detergent in the protein extraction buffer; since NOD is an integral plasma membrane protein, we expected that NOD might interact with other plasma membrane proteins. Separately, we used NOD as bait for a quantitative yeast two-hybrid screen against a prey library cloned from normalized maize cDNA. This experiment was conducted in parallel alongside several unrelated baits, which allowed us to filter any recurring hits as false positives.

**Figure 1.**
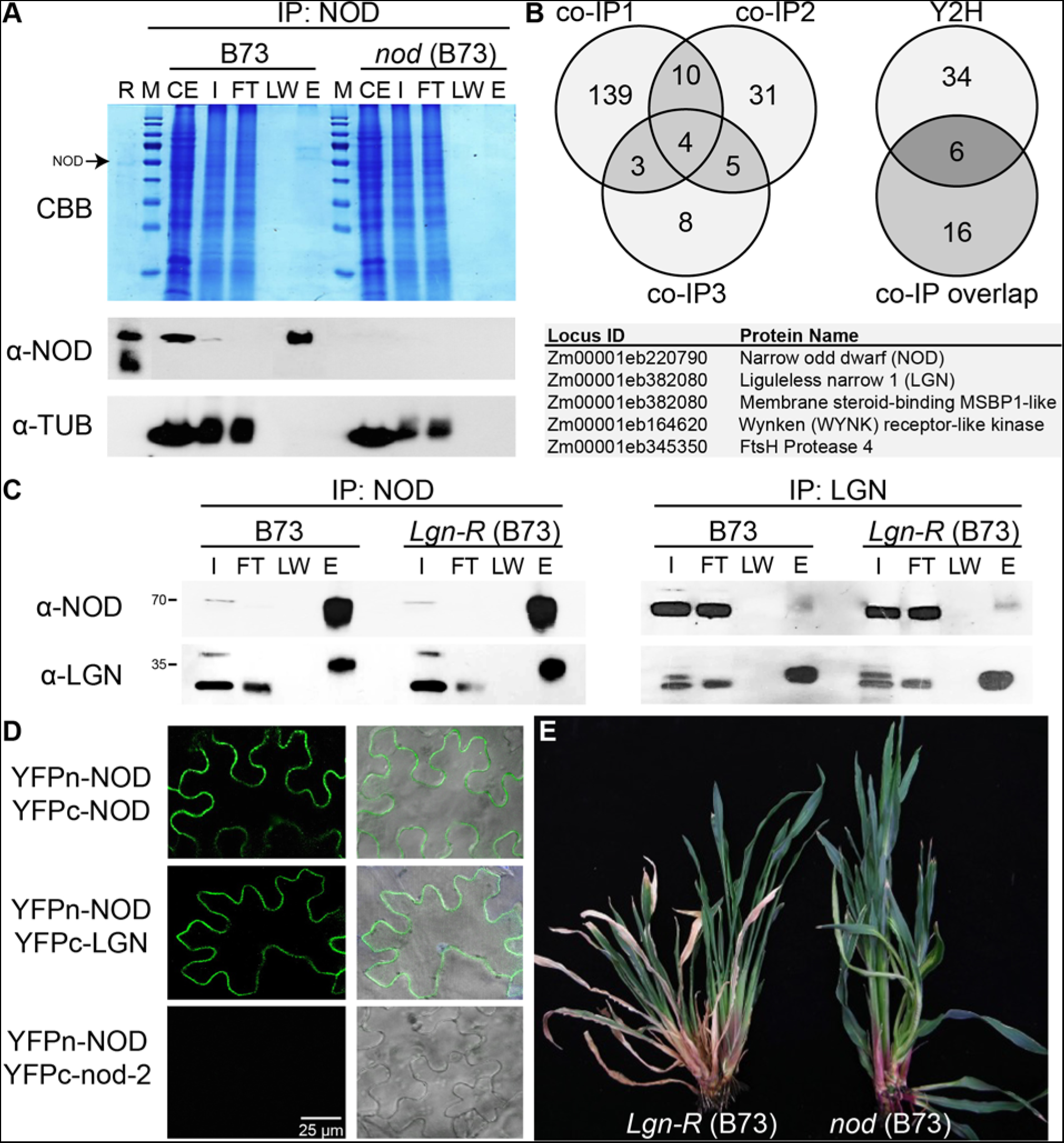
LGN and NOD proteins can interact and cause similar pleiotropic phenotypes. **(A)** To identify interacting proteins, NOD was immunoprecipitated using a native polyclonal antibody from wild-type B73 seedlings and null *nod* mutant B73 seedlings. As shown here, NOD was strongly enriched in the eluate after immunoprecipitation. Lanes are labeled to indicate crude extract (CE), input (I), flow-through (FT), last wash (LW), and eluate (E). **(B)** Coimmunoprecipitates of NOD were identified by mass spectrometry in three experiments. 200 potential interactors were identified, with several overlapping candidates between at least two experiments. A yeast two-hybrid (Y2H) screen for NOD interactors identified 40 potential interactors, including 6 of the candidate interactors that had been found in at least two co-IP experiments. Select interactors of interest are shown here, including several receptor-like proteins. **(C)** To validate the putative interaction between LGN and NOD, reciprocal co-immunoprecipitation experiments were conducted using native polyclonal antibodies raised against each protein. Both experiments confirmed that LGN and NOD can co-immunoprecipitate with each other, supporting the hypothesis that they physically associate *in vivo*. **(C)** Bimolecular fluorescence complementation assays were conducted by heterologously expressing NOD, LGN, or nod-2 protein that mislocalizes to cytoplasmic aggregates tagged with split-YFP constructs in *Nicotiana benthamiana* leaves (the N-terminal portion of YFP is indicated as YFPn, the C-terminal portion of YFP is indicated as YFPc). Epidermal fluorescence at the plasma membrane confirms that NOD can interact with itself and that NOD can interact with LGN. As expected, mis-localized nod-2 protein cannot interact with NOD, which serves as a negative control. **(D)** Photo of *Lgn-R/Lgn-R* and *nod/nod* mutants in the B73 background. Both mutants display pleiotropic defects in growth and development, including arrest of the primary shoot, excessive tillering, short and narrow leaves, and absence of a proper ligule.

Between the Y2H and coIP experiments, we identified over 200 potential interactors of NOD (Fig. 1B). These included NOD itself, as expected, since the Arabidopsis orthologue of NOD forms homomultimeric complexes (Yoshimura, Iida and Iida, 2021); the NOD-NOD interaction was further validated using bimolecular fluorescence complementation after tagging NOD with split YFP constructs and expressing the proteins heterologously in *Nicotiana benthamiana* epidermal cells (Fig. 1D). We also identified several plasma membrane-localized proteins interacting with NOD, including Liguleless narrow 1 (LGN), a BRASSINOSTEROID SIGNALING KINASE-like kinase that we named WYNKEN for its association with NOD, and a membrane steroid-binding protein (MSBP1)-like protein. Several interactors were unexpected, including, e.g., the orthologue of Arabidopsis FtsH4, a mitochondrial protease. Given that FtsH4 is targeted to mitochondria, this could be an artifactual interaction that does not occur *in vivo*, but FtsH4 was identified in both coIP and Y2H screens, suggesting that this candidate could warrant future investigation (Fig. 1B). Among the candidate interactors, we were most intrigued by the putative association with LGN, given that *Lgn-R* mutants show similar phenotypes to *nod* mutants (Fig. 1E), including stunted growth, defective ligule development, and constitutive induction of biotic stress responses. Therefore, we hypothesized that these proteins could act in overlapping or epistatic pathways, and decided to focus on defining the potential relationship between LGN and NOD.

To validate the LGN-NOD interaction, we first performed reciprocal coIP experiments using polyclonal antibodies against NOD and LGN to precipitate one of the proteins, followed by probing eluates with Western blots. As predicted, LGN was detected in the α-NOD immunoprecipitates (Fig. 1C, left panel) and NOD was detected in the α-LGN immunoprecipitates (Fig. 1C, right panel). Similar results were obtained in *Lgn-R* mutants, where the LGN-R protein is stably expressed but is catalytically dead, revealing that the NOD-LGN interaction is likely independent of LGN’s kinase activity. BiFC also confirmed that NOD and LGN can interact with each other, and that this interaction occurs in the plasma membrane (Fig. 1D). Therefore, using several different experimental approaches, we confirmed that NOD and LGN can associate with each other in plant cells.

### LGN is capable of phosphorylating NOD

Since LGN is a receptor-like kinase, we sought to test whether LGN might phosphorylate NOD (Fig. 2). Recombinant LGN and NOD proteins were co-incubated in kinase reaction buffer alongside various controls, including single protein incubations and co-incubation with the kinase-dead LGN-R protein. Protein phosphorylation was first analyzed by SDS-PAGE followed by staining with either Pro-Q Diamond, a stain that specifically detects phosphorylated proteins, or Coomassie Brilliant Blue, which indiscriminately stains all proteins (Fig. 2A). As previously shown, LGN autophosphorylates (resulting in slower migration during SDS-PAGE and staining with Pro-Q Diamond), whereas LGN-R does not autophosphorylate (and therefore migrates more quickly during SDS-PAGE and is not detected by Pro-Q Diamond stain) (Fig. 2A). NOD is readily phosphorylated by LGN *in vitro* (Fig. 2A), but not when it is incubated by itself or with LGN-R, confirming that the phosphorylation observed in this assay is LGN-dependent. To validate these results, we analyzed putative NOD and LGN phosphoproteins by mass spectrometry (Fig. 2B, 2C). These analyses revealed at least 15 possible autophosphorylated sites in LGN and at least 7 potential LGN-catalyzed phosphorylation sites in NOD. Unexpectedly, these included serine, threonine, and tyrosine residues, which indicates that LGN is capable of acting as a dual-specificity Ser/Thr and Tyr kinase, at least under the *in vitro* conditions used in this experiment.

**Figure 2.**
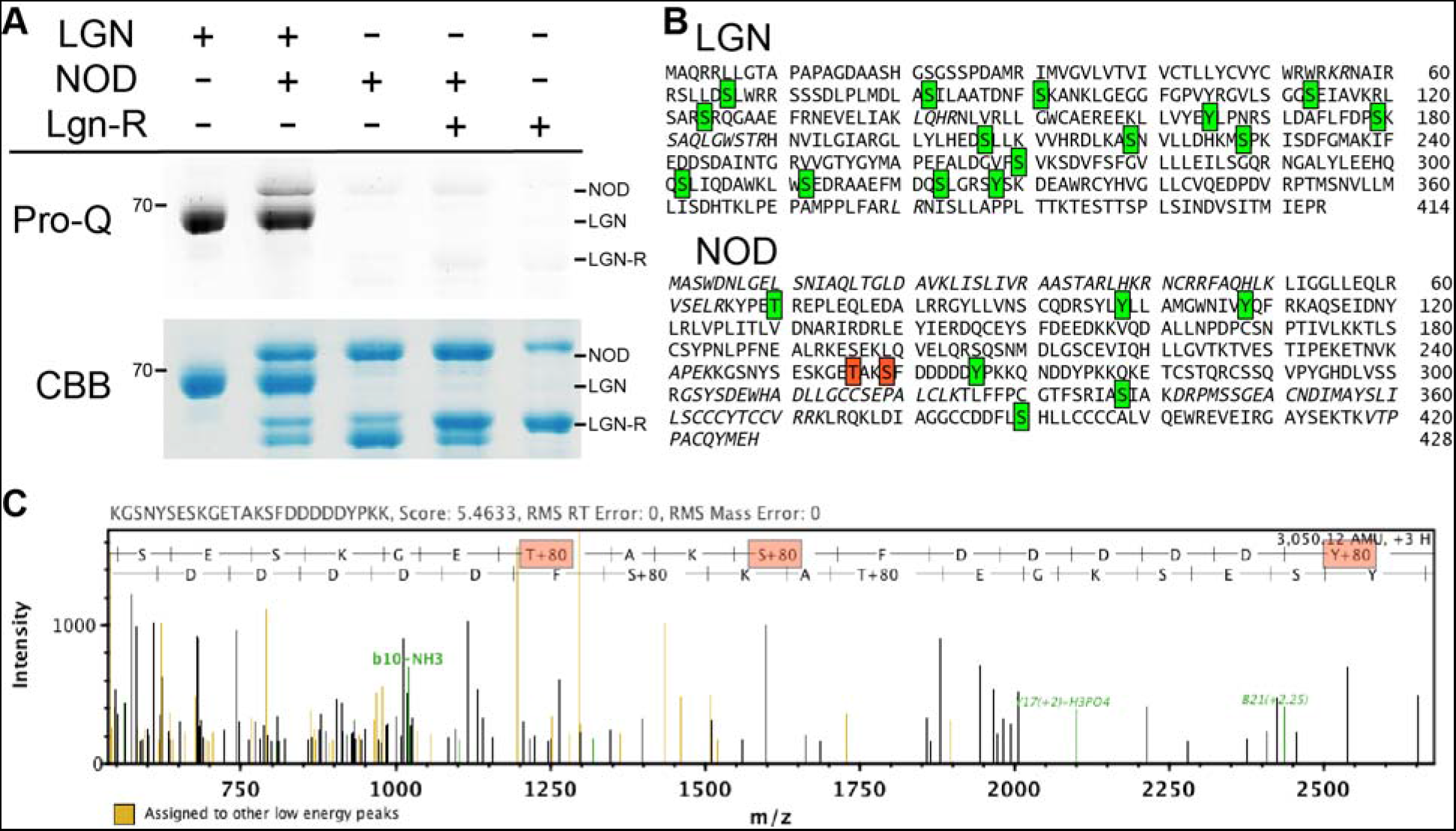
LGN can phosphorylate NOD *in vitro*. **(A)** Recombinant LGN, NOD, and catalytically-inactive LGN-R proteins were incubated as indicated in kinase buffer. The proteins were then separated by SDS-PAGE and stained with either Pro-Q Diamond (Pro-Q), which stains phosphoproteins, or Coomassie Brilliant Blue (CBB), which stains total proteins. LGN can phosphorylate itself, as previously shown, and can also phosphorylate NOD. Negative controls confirm that NOD was not phosphorylated in the absence of functional LGN enzyme. **(B)** After incubation as in panel A, proteins were analyzed by mass spectrometry, which identified multiple putative phosphosites (green boxes, high-confidence; red boxes, could not distinguish) on both LGN and NOD. **(C)** Representative spectrum from mass spectrometry analysis of recombinant proteins after co-incubating in kinase buffer.

### Developmental defects are more severe in the double *nod;Lgn-R/+* mutants than either single mutant

Building on the compelling evidence that LGN and NOD molecularly interact in maize cells and that *Lgn-R* and *nod* mutants share similar phenotypes, we hypothesized that *Lgn-R* and *nod* mutants might genetically interact. The phenotype of each single mutant varies considerably in different inbred backgrounds, so we made crosses of *nod* to *Lgn-R* that had both been introgressed into two inbred backgrounds, A619 and B73. In both inbreds, *Lgn-R* heterozygotes were crossed to *nod* mutants and back-crossed to *nod* to generate families that segregated single mutants, double mutant, and non-mutant siblings (see methods for crossing details).

In both A619 and B73 backgrounds, we consistently observed that the *nod;Lgn-R*/+ double mutants exhibited much more severe defects in growth and development than either single mutant. For example, in B73, the blades of leaf 10 are somewhat shorter and ~50% narrower in *Lgn-R/+* plants, yet shorter and narrower in *nod* plants, but shortest and narrowest in the *nod;Lgn-R/+* double mutants, compared to wild-type plants grown under the same conditions (Fig. 3A, 3E). Similarly, in A619, *Lgn-R*/+ plants are ~20% shorter with leaves ~15% narrower than wild-type and *nod* plants are ~70% shorter with leaves ~60% narrower than wild-type (Fig. 3B, 3C). Consistently, *nod;Lgn-R*/+ plants are ~90% shorter and have ~85% narrower leaves than wild-type, and are therefore much smaller than either single mutant (Fig. 3B, 3C). Focusing on developmental patterning at the blade-sheath boundary, *nod;Lgn-R/+* is again more severe than either single mutant. The ligule is somewhat reduced in heterozygous *Lgn-R/+* plants and only present near the leaf margins in *nod* plants, but virtually absent in *nod;Lgn-R/+* mutants (Fig. 3D, upper and middle panels). Similarly, the remaining hints of an auricle and blade-sheath boundary in *nod* mutants are both completely abolished in *nod;Lgn-R/+* mutants (Fig. 3D, middle and lower panels). The main shoots of both *nod* and *nod;Lgn-R/+* mutants die and the plants are highly tillered (producing 2 ± 0.4 tillers on average, with no significant difference in tiller number between genotypes, *p* = 0.84 by Student’s *t* test, *n* ≥ 8), although *nod* tillers are significantly longer than *nod;Lgn-R/+* tillers (19.9 ± 1.9 cm for *nod* versus 6.7 ± 0.6 cm for *nod;Lgn-R/+*, *p* < 10^−6^ by Student’s *t* test, *n* ≥ 15). Based on these morphological results, we concluded that *nod* and *Lgn-R/+* are not epistatic with each other, but instead have cumulative effects on maize growth and development.

**Figure 3.**
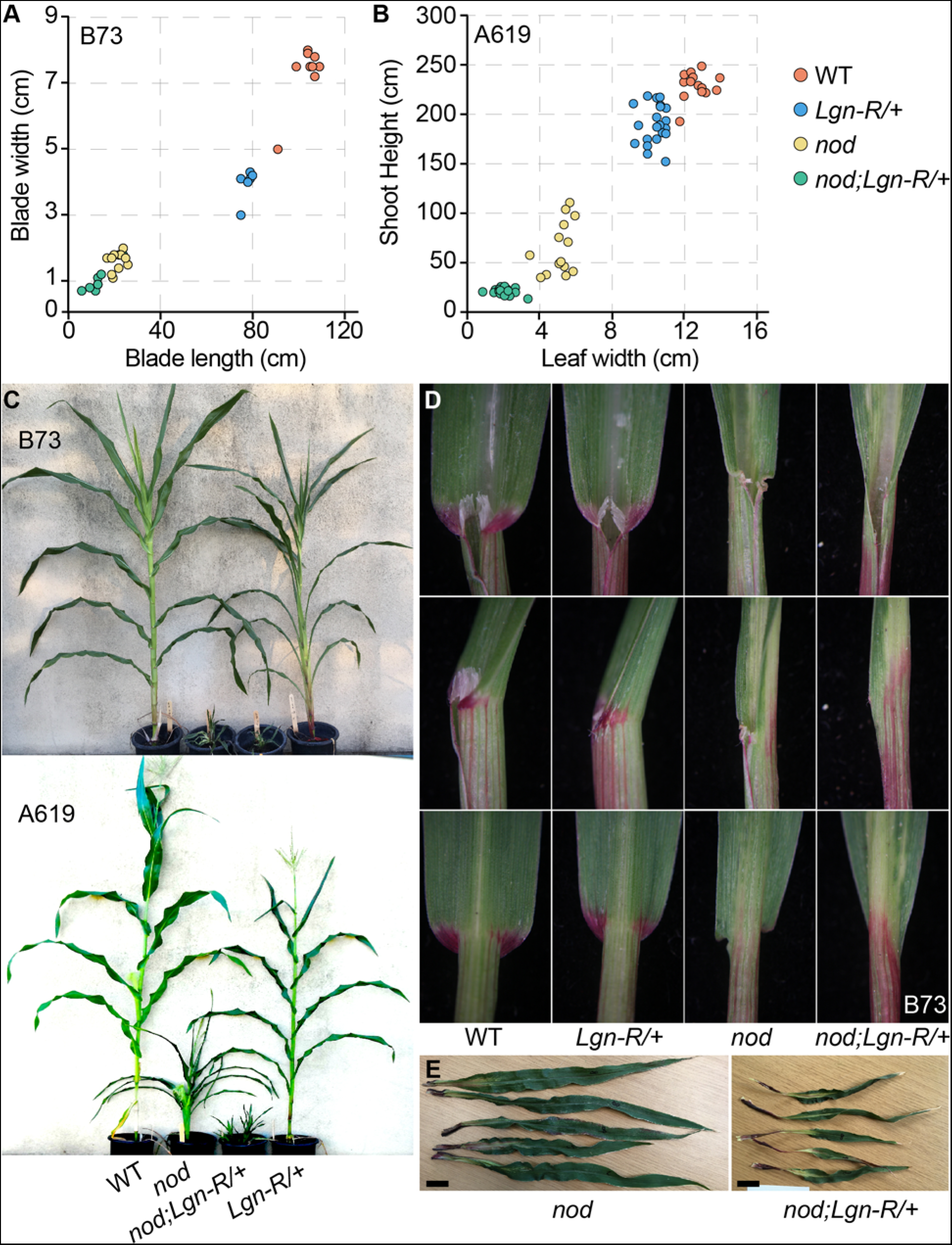
*nod;Lgn-R/+* double mutants exhibit more severe phenotypes than either single mutant in both A619 and B73 inbred backgrounds. **(A)** Blade width and length of leaf 8 were measured for all four genotypes in B73. The genotypes clearly separated into four groups, with the most severe growth defects in the *nod;Lgn-R/+* double mutants. **(B)** Plant height and leaf width were measured for all four genotypes in A619 at maturity. The genotypes clearly separated into four groups with apparently additive effects on these traits in the *nod;Lgn-R/+* double mutant. **(C)** Photos of each genotype grown in the greenhouse. The B73 family was photographed prior to maturity for comparison to the severe genotypes. The A619 family was grown to maturity. **(D)** Representative photos of leaf 2 for all four genotypes in B73. *Lgn-R/+* heterozyogtes have somewhat reduced ligules and *nod* homozygotes have severely reduced ligules, but the double *nod;Lgn-R/+* mutants have almost no visible ligule (top and middle panels). Moreover, auricles and any other indications of a defined blade-sheath boundary are somewhat disrupted in *nod* single mutants but virtually absent in the double mutants (bottom panel). **(E)** Leaf 8 of *nod* or *nod;Lgn-R/+* mutants in the B73 background were photographed to highlight the enhanced growth defects in double mutants. Images are at the same scale.

### NOD and LGN are required to repress a shared stress-response transcriptional network

To better understand the extreme phenotypes of *nod;Lgn-R*/+ double mutants, we used global profiling methods to define the effects of the single and double mutants on the maize transcriptome and metabolome in both B73 and A619 (Supplementary Dataset 1). Metabolomics was carried out on the second leaf blade of plants at V4 and the shoot apex was used for RNAseq (see methods).

At the transcriptomic level in both A619 and B73 backgrounds, we observed significant overlap among all three mutant genotypes (relative to wild-type; a significantly differentially expressed gene, which we abbreviate DEG, showed a difference in expression relative to wild-type negative controls with *p_adj_* < 0.05, adjusted to reduce false positive results). The most striking difference in transcriptomes was a background-dependent genetic interaction with the single mutants: in B73, *nod* caused much more drastic transcriptional reprogramming, with almost four-fold more DEGs in *nod* plants (~7,300 DEGs) than *Lgn-R/+* plants (~1,900 DEGs) (Fig. 4A). Conversely, *Lgn-R/+* had a stronger effect on the transcriptome in A619, with more than five-fold more DEGs in *Lgn-R/+* plants (~6,600) than in *nod* plants (~1,200 DEGs) (Fig. 4D). This result was initially surprising, since *Lgn-R/+* plants show more potent defects in growth and development in B73 than in A619. As discussed in detail below, however, these transcriptomic results are supported by the mutant metabolomes, which show the same relationship: *nod* is more metabolically disruptive in B73 and *Lgn-R/+* is more metabolically disruptive in A619.

**Figure 4.**
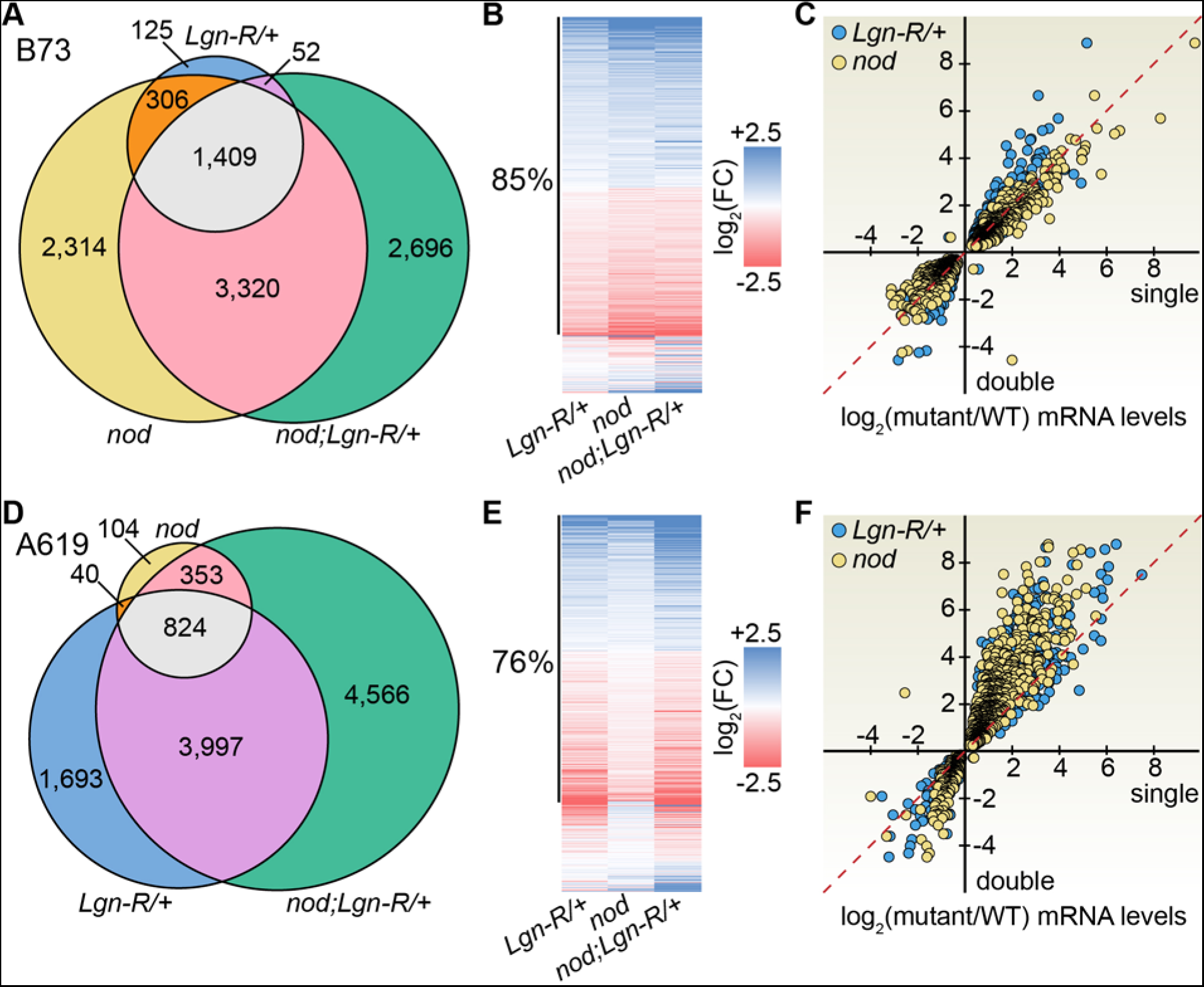
RNA-Seq reveals overlapping effects of *Lgn-R/+*, *nod*, and *nod;Lgn-R/+* mutants on transcriptional programs in B73 and A619 backgrounds. **(A)** Significantly differentially-expressed genes (DEGs, *p_adj_* < 0.05) detected in one, two, or all three genotypes in the B73 background. Note that almost all *Lgn-R/+* DEGs were also detected in *nod*. **(B)** Heatmap showing changes in mRNA level (relative to wild-type) for any gene detected as significantly affected in at least one genotype in the B73 background (i.e., all DEGs in panel A). Although many DEGs met stringent statistical significance thresholds only in *nod* or *nod;Lgn-R/+*, 85% of DEGs were coordinately impacted across all genotypes. **(C)** Scatterplot showing the magnitude of change in gene expression for the 1,409 DEGs significantly affected in all three genotypes in the B73 background, shown as logarithm of fold-change in expression (relative to wild-type) for single mutants (*x*-axis) versus the double mutant (*y*-axis). Almost none of these DEGs are oppositely affected in the genotypes, and the magnitude of effects is similar among all three genotypes (the slope of regression lines is ~1.0, which is shown with a red dashed line). **(D)** DEGs detected in one, two, or all three genotypes in the A619 background. Note that most *nod* DEGs were also detected in *Lgn-R/+*. **(E)** Heatmap as in panel B, but for the A619 background. Although many DEGs met stringent statistical significance thresholds only in *Lgn-R/+* or *nod;Lgn-R/+*, 76% of DEGs were coordinately impacted across all genotypes. **(F)** Scatterplot as in panel C, but for the 824 DEGs affected in all mutants in the A619 background. Virtually all of these DEGs are coordinately impacted across the three genotypes. The magnitude of change in gene expression is somewhat greater in the double mutant (the slope of linear regression lines is 1.4 for *Lgn-R/+* and 1.6 for *nod*, with *r^2^* > 0.85; a red dashed line shows what a slope of 1.0 would look like).

Although the sheer number of DEGs differed among the mutant genotypes, the vast majority of DEGs overlapped between the *nod* and *Lgn-R/+* mutants, and virtually all of these DEGs were coordinately impacted (induced in all mutant genotypes). In B73, 1,715 (91%) of the 1,892 DEGs detected in *Lgn-R/+* were also detected as DEGs in *nod* (Fig. 4A), and only 1 of these was oppositely impacted (induced in at least one mutant genotype and repressed in at least one other). To phrase this more directly, nearly all of the transcriptomic changes caused by *Lgn-R/+* were also caused by *nod* in the B73 background. Predictably, the vast majority of overlapping DEGs in both single mutants were also coordinately impacted DEGs in the double mutant. The same overall pattern holds true in A619: 65% of DEGs in *nod* overlapped with *Lgn-R/+* (864 / 1,321 DEGs), and 99.9% of these DEGs were coordinately impacted (863 / 864 DEGs). Over 95% of the overlapping DEGs between the single mutants were also detected in the double mutant, and, again, virtually all changes in gene expression were coordinate among the three genotypes. Therefore, at a transcriptional level, *Lgn-R/+* and *nod* act on almost fully overlapping pathways, although one mutant has a stronger overall impact on the maize transcriptome than the other depending on the genetic background.

We next focused on determining whether simultaneously disrupting LGN and NOD caused any transcriptomic effects that could not have been predicted from the single mutant transcriptomes (i.e., signatures of epistasis or synergy). Indeed, many more DEGs were detected in the double mutants than in either single mutant background, which was not surprising, given their more severe phenotypes. We observed no clear cases of discord among the transcriptomes, however, with nearly all of the overlapping DEGs between the double mutants and either single mutant were coordinately induced or coordinately repressed (Fig. 4B,E). In fact, if we ignored cutoffs for statistical significance and simple compared the change in gene expression for any gene detected as a DEG in one of the three mutant genotypes, almost all genes were still coordinately impacted (85% in B73 and 76% in A619, shown as clusters in Fig. 4B and Fig. 4E, respectively). Thus, while there are certainly differences in mRNA levels among mutant genotypes in a given inbred background, the overall effects of *nod*, *Lgn-R/+*, or their combination in *nod;Lgn-R/+* are extremely similar, clearly supporting the hypothesis that LGN and NOD act through comparable signal transduction pathways to coordinate nuclear gene expression.

Quantitatively, the DEGs overlapping among the single and double mutants generally showed a greater magnitude of differential expression in the double mutant in A619, where the double mutant had a much stronger growth and development phenotype than either single mutant. On average, in A619 the log_2_(fold-changes in mRNA levels) in the double mutant were ~1.4x greater in magnitude than in *Lgn-R/+* and ~1.6x greater in magnitude than in *nod* (Fig. 4F). Thus, although the double mutant often coordinately impacted the expression of the same genes as the single mutants, the degree of differential expression showed some cumulative enhancement in the double mutants. We did not observe a similar cumulative effect in B73 (Fig. 4C), perhaps because the *nod* phenotype is already so severe in the B73 background that the changes in gene expression are effectively already “saturated” in the single mutants. This hypothesis is corroborated by the already very severe developmental phenotypes in *nod* single mutants.

Lastly, we analyzed the mutant transcriptomes using MapMan gene ontologies to identify the biological processes regulated by *nod*, *Lgn-R/+*, and the double mutants at the transcriptional level (Fig. 5). Statistical tests for enrichment of MapMan gene ontologies revealed transcriptional induction of biotic stress responses and secondary/specialized metabolic pathways across the mutant genotypes, whereas genes involved in cell division and cell cycle progression are broadly repressed in the mutants (Fig. 5A). These effects are similar to previously described transcriptional responses to *Lgn-R* and *nod* in B73 (Rosa *et al*., 2017; Anderson *et al*., 2019). Given the scale of transcriptional reprogramming discovered in each mutant genotype, we focused especially on significantly induced categories of transcription factors that could mediate responses to *nod* and/or *Lgn-R/+*. Diverse genes of the family of APETALA2/ETHYLENE RESPONSE FACTOR (AP2/ERF)-like transcription factors, which often promote the expression of stress response genes in plants, are significantly transcriptionally induced in *nod*, *Lgn-R/+*, and *nod;Lgn-R/+* in the B73 background (Fig. 5B). Moreover, in both A619 and B73 backgrounds, mRNAs of genes that encode dozens of NAC- and WRKY-type transcription factors are also induced; NAC and WRKY transcription factors are also established mediators of abiotic and biotic stress responses in plants (Fig. 5B) (Yu, Chen and Chen, 2001; Rushton *et al*., 2010; Puranik *et al*., 2012; Fu and Dong, 2013; Nuruzzaman, Sharoni and Kikuchi, 2013; Shao, Wang and Tang, 2015; Phukan, Jeena and Shukla, 2016; Viana *et al*., 2018). Overall, these transcriptional patterns illustrate the concept of “growth-defense tradeoffs”, with enhanced expression of genes involved in defense against pathogens and abiotic stress coupled with repressed expression of genes involved in cell division and growth (Fig. 5C).

**Figure 5.**
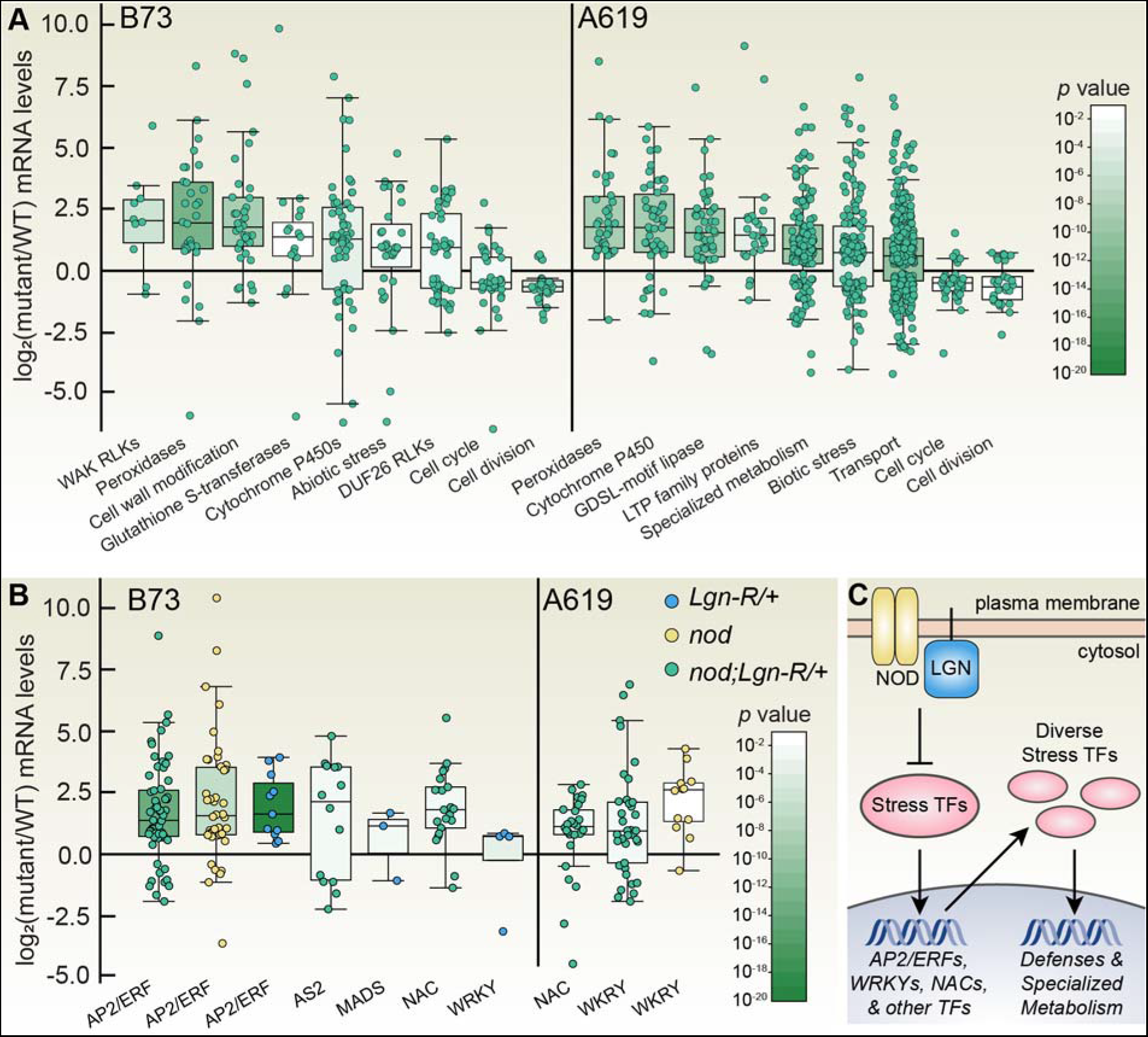
Gene ontology analysis of biological processes impacted at the transcriptional level in *nod*, *Lgn-R/+*, and *nod;Lgn-R/+* mutants. **(A)** Select significantly impacted processes identified by MapMan in the double mutants are shown, with *p-*value for the category (as determined by a Mann-Whitney test with the Benjamini-Yekutieli procedure to correct for false positives) indicated by shade of green, box-whisker plots drawn with Tukey’s method showing the logarithm of the change in mRNA level for genes (relative to wild-type) in the indicated category, and arranged in order by median change in gene expression for the category. Genes involved in stress signaling and responses were induced in the mutants, such as receptor-like kinases (RLKs), redox stress-related peroxidases and glutathione S-transferases, lipid transfer proteins (LTPs), stress-induced GDSL-type lipases, and categories involved in specialized/secondary metabolism (including cytochrome p450 enzymes and transporters). In contrast, genes involved in cell cycle progression and cell division were largely repressed in the mutants. **(B)** Significantly induced categories of transcription factors (TFs) shown as box-whisker plots (as in panel A) for all three mutant genotypes in both inbreds. TF families strongly associated with promoting stress responses, including AP2/ERF, NAC, and WRKY TFs, were strongly induced across genotypes. **(C)** Cartoon suggesting that loss of cell-surface proteins LGN and/or NOD de-represses stress response transcriptional networks (as seen in panel A) that is mediated, in part, by the potent transcriptional induction of stress responses TFs (as seen in panel B).

### *nod* and *Lgn-R* cause similar disruptions to primary and secondary metabolism

Metabolomic analysis supported the hypothesis that *nod* and *Lgn-R* act on overlapping pathways that are most strongly affected in the double mutant. Principal components analysis of the metabolomes showed that the largest principal components (PC1: 38.1% of variation in B73, 38.9% of variation in A619, Fig. 6A,B) in both A619 and B73 were sufficient to separate wild-type, *nod*, *Lgn-R/+*, and double mutants from each other in a linear “spectrum.” Correlating with the order of phenotypic severity, in B73, the spectrum of PC1 showed that *Lgn-R/+* is most metabolically similar to wild-type, followed by *nod*, and finally the double mutant, whereas in A619, the spectrum of PC1 showed that *nod* is most metabolically similar to wild-type, followed by *Lgn-R/+*, and finally the double mutant. The second principal components (PC2: 15.7% of variation in B73, 11.8% of variation in A619, Fig. 6A, 6B) could not be used to separate the genotypes, however, indicating that PC2 reflects individual-to-individual variation independent of *nod* and *Lgn-R*. Overall, this clear result suggests that *nod* and *Lgn-R* broadly impact the same metabolic processes (represented here by PC1) and that the double mutant cumulatively exacerbates these metabolic impacts.

**Figure 6.**
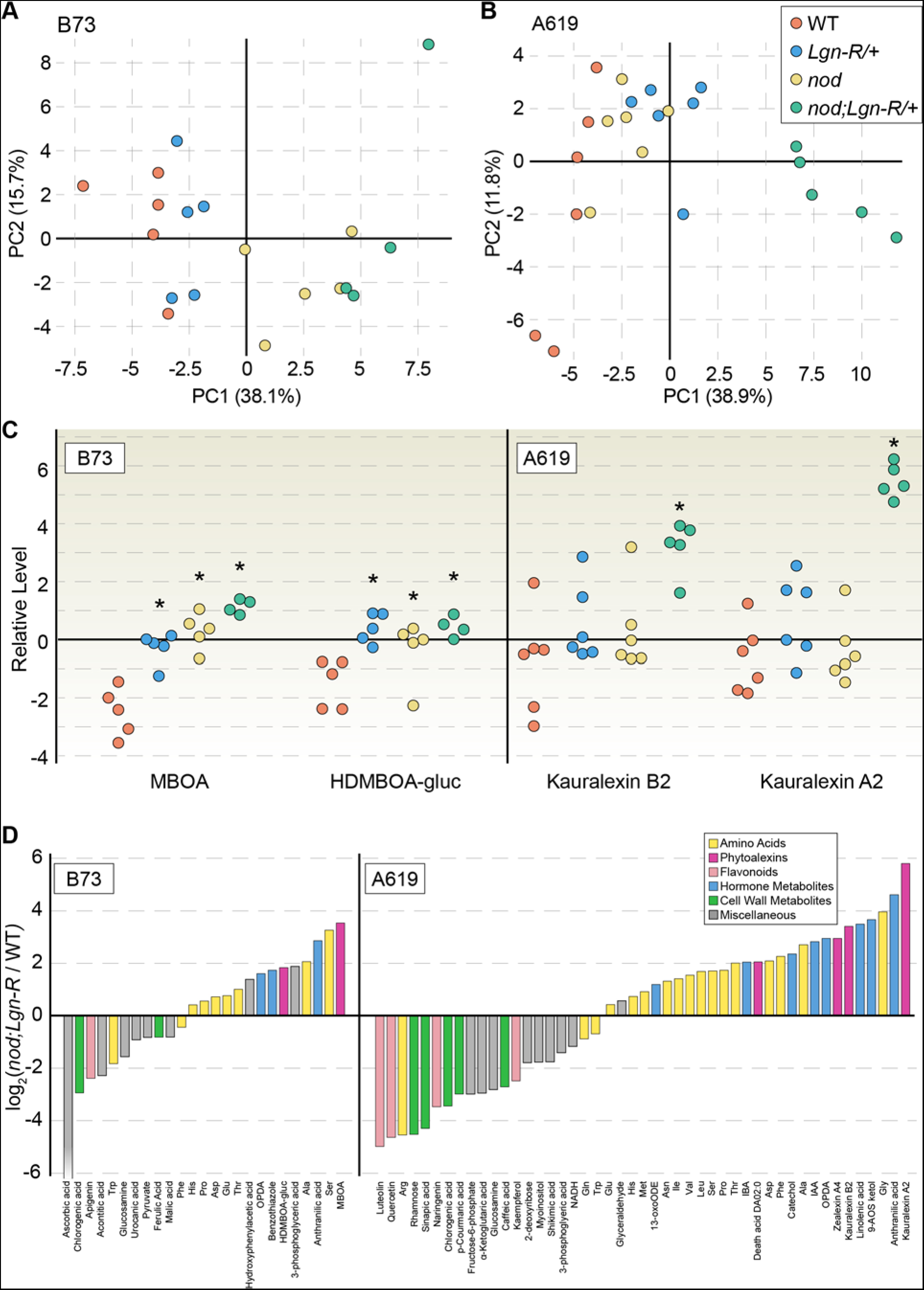
Metabolomic analyses confirm that NOD and LGN act in overlapping pathways to coordinate plant physiology and development. **(A,B)** Principal components analysis of untargeted metabolomic analyses of wild-type (red), *Lgn-R/+* (blue), *nod* (yellow), and *nod;Lgn-R/+* (green) seedling shoots in either B73 (A) or A619 (B) genetic backgrounds revealed that a single principal component, PC1, accounts for over 38% of variation among samples. PC1 can be used to distinguish all four genotypes along a linear “spectrum” of phenotypic severity, with the double mutant representing the cumulative effects of *nod* and *Lgn-R/+* on overlapping metabolic pathways. **(C)** Phytoalexins overaccumulate in all mutants, with the highest levels observed in *nod;Lgn-R/+* double mutants. The class of phytoalexins induced was different between the two genetic backgrounds: in B73, benzoxazinoids (MBOA, HDMBOA-glucose) were induced, whereas in A619, terpenoids (kauralexins) were induced. **(D)** Significantly affected metabolites (*p* < 0.10, Student’s *t* test) quantified in the untargeted metabolomic analyses of the double mutant *nod;Lgn-R/+* compared to wild-type are shown here, color-coded by various categories. In both backgrounds, free amino acids, phytoalexins, and phytohormones generally accumulated to higher levels in the mutants, whereas flavonoids, cell wall metabolites, and primary metabolites accumulated to lower levels in the mutants.

Upon closer examination of the many compounds that differentially accumulate in the mutants, a consensus pattern emerges: in *nod*, *Lgn-R/+*, and *nod;Lgn-R/+* mutants, metabolites involved in biotic stress responses are induced while metabolites involved in growth and development, such as free amino acids, cell wall synthesis intermediates, and growth-regulating phytohormones, are dysregulated. Phytoalexins, including terpenoids (e.g., kauralexins and zealexins) and benzoxazinoids (e.g., 6-methoxybenzoxazolinone or MBOA), are strongly induced across all genotypes (Fig. 6C). Phytoalexins are secondary metabolites with antimicrobial properties that are most often synthesized upon pathogen attack as a defense response, but are rarely detected at significant levels in healthy plants (Ahuja, Kissen and Bones, 2012). Although phytoalexins were strongly induced in all mutant genotypes, different families of phytoalexins were induced depending on the genetic background, perhaps reflecting differences in immune signaling and/or metabolic pathways in these distinct inbred lines. In B73, benzoxazinoids are induced in the single and double mutants, with slightly stronger induction in the double mutants. In A619, terpenoids are instead somewhat induced in all mutants, although the most significant and drastic effect is only observed in the *nod;Lgn-R/+* double mutant.

Since the principal components analysis confirmed that the double mutants represent the cumulative effects observed in both *nod* and *Lgn-R/+* single mutants, here, we will focus our detailed metabolomic analysis on the double mutants in the A619 and B73 backgrounds. Dozens of metabolites accumulated to significantly different levels in the mutants (Fig. 6D). The most extreme example is ascorbic acid: in the B73 background, ascorbic acid levels were ~2-fold decreased in *Lgn-R/+* mutants, but barely detected at all in *nod* or *nod;Lgn-R/+* mutants. Ascorbic acid (vitamin C) scavenges reactive oxygen species (ROS) to limit redox stress; the low levels of ascorbic acid in these mutants correlates with the observed induction of redox stress and ROS levels previously described in *nod* leaves (Rosa *et al*., 2017). Hormone metabolism is disrupted (Fig. 6D, blue bars), including overaccumulation of auxins, oxylipins (the class of hormones that includes jasmonic acid), cytokinin-like benzothiazole, and salicylic acid-related compounds (precursor anthranilic acid and catabolism product catechol). Overaccumulation of any of these compounds on their own would likely disrupt plant development and physiology, so the coordinated induction of these compounds in the *nod;Lgn-R/+* plants likely contributes to their pleiotropic phenotypes.

Most free amino acid levels are significantly higher in the mutants (Fig. 6D, yellow bars), indicating that protein synthesis is inefficient, which would correlate with the extreme *nod;Lgn-R/+* growth defects. Similarly, the extremely low levels of cell wall synthesis intermediates, including the chlorogenic acids required for lignin biosynthesis, may also correlate with reduced cell wall expansion (i.e., reduced growth) and disrupted developmental patterning (Fig. 6D, green bars). Indeed, phloroglucinol staining of leaf and stem cross-sections in *nod* mutants previously revealed the absence of differentiated lignified cells that were present in wild-type stems and leaves (Rosa *et al*., 2017). Lastly, many intermediates in primary metabolism, including citric acid cycle and glycolysis intermediates, are significantly downregulated in the double mutants (Fig. 6D, gray bars), confirming that the severe growth defects in *nod;Lgn-R/+* plants correlate with the disruption of the core metabolic pathways.

Finally, we hypothesized that some of the metabolic changes in the mutants could be due to specific transcriptional changes in gene expression. Metabolism is often controlled not only at the transcriptional level, but also by post-transcriptional regulation of gene expression and various biochemical feedback mechanisms acting on enzymes, so core metabolite levels are necessarily expected to correlate with enzyme-encoding mRNA levels. Pathogen and herbivore defense metabolites, however, are often synthesized by highly specialized pathways that are transcriptionally induced only under certain circumstances, e.g., upon recognition of a pathogen infection. Here, for example, we did observe significant, strong transcriptional induction of genes encoding enzymes that promote terpenoid synthesis, including *TERPENE SYNTHASE*. In contrast to the coordinated induction of terpenoid synthesis genes and accumulation of terpenoid phytoalexins in the mutants, not every secondary metabolic change correlated with the transcriptome. Consistently, in all mutants and both backgrounds, flavonoid levels were significantly lower (Fig. 6D). *FLAVONOL SYNTHASE* transcripts were significantly induced in the mutants, however. These results illustrate that enzyme transcript levels do not always positively correlate to the level of their chemical products, as has been demonstrated repeatedly in other conditions and biological systems (Fernie and Stitt, 2012; Melandri *et al*., 2022).

## DISCUSSION

Here, we demonstrated that two maize proteins localized to the plasma membrane, the putative mechanosensitive Ca^2+^ channel NOD and the receptor-like kinase LGN, physically associate with each other at the molecular level and coordinate largely overlapping signaling networks with striking effects on maize development and physiology. Genetic disruption of either NOD or LGN activity in the null *nod* mutants or semi-dominant negative (antimorph) *Lgn-R/+* mutants is sufficient to disrupt growth, but complete loss of both gene functions in double *nod;Lgn-R/+* mutants causes even more severe phenotypes. Using unbiased profiling approaches to define molecular phenotypes of the single and double mutants, we found that all three genotypes reprogram maize biology to promote stress and pathogen defense responses at the cost of reducing resources allocated to growth and development, an illustration of the “growth-defense trade-off” hypothesis.

The precise molecular function of NOD remains unclear, although its downstream effects are consistent and reproducible: NOD is required to suppress a stress responsive nuclear transcriptional program. NOD encodes a PLAC8 domain and is structurally related to genes including *CELL NUMBER REGULATOR 1 (CNR1)* in maize and *FW2.2*, a gene responsible for a major quantitative trait locus for fruit weight in tomato (Frary *et al*., 2000; Guo *et al*., 2010). Moreover, a recent study found that some polymorphisms in *NOD* alleles among maize genotypes are significantly associated with yield-related traits in ear morphology (Zuo *et al*., 2021). Thus, NOD and NOD-like proteins play crucial roles in agricultural species, and they could even be useful targets for plant breeders (Beauchet *et al*., 2021). Mechanistically, the Arabidopsis orthologues of NOD were first identified for their ability to complement loss of a Ca^2+^ channel in yeast (and thus named MID1-COMPLEMENTING ACTIVITY, or MCA), hinting that NOD could act as a plasma membrane Ca^2+^ channel (Nakagawa *et al*., 2007), although there have been conflicting reports using different methodologies to test whether NOD is capable of trafficking ions across plasma membranes in heterologous *Xenopus* oocyte systems (Furuichi *et al*., 2012; Rosa *et al*., 2017). Regardless, this study reveals that NOD acts on pathways that substantially overlap with the LGN-regulated pathways in maize cells, implying that studies of the cellular networks downstream of both NOD and LGN may be more biologically informative than continued debates on the precise molecular activity of the NOD protein.

*nod* and *Lgn-R* mutants both show strong background-dependent phenotypes. In both cases, the phenotypes are most severe in the B73 background, moderate in the A619 background, and mild in the Mo17 background. Quantitative trait association mapping showed that a single locus, *Sympathy for the Ligule (Sol)*, is largely responsible for the background-dependent defects in *Lgn-R* between B73 and Mo17 (Buescher *et al*., 2014). *Sol* encodes an orthologue of Arabidopsis ENHANCED DISEASE RESISTANCE 4 (EDR4), a protein that represses mitogen-activated protein (MAP) kinase signaling networks through molecular interactions with the upstream MAP3K (MEKK) ENHANCED DISEASE RESISTANCE 1 (EDR1) (Wu *et al*., 2015; Anderson *et al*., 2019). Phosphoproteomic analysis of *Lgn-R/+* mutants hinted that Lgn might function in a common receptor-like kinase role by repressing a MAP kinase cascade that activates stress-responsive WRKY transcription factors, and that the Mo17 allele of *Sol* is capable of partially repressing that MAP kinase cascade, whereas the B73 allele of *Sol* cannot (Anderson *et al*., 2019). The genetic modifiers responsible for background-dependent phenotypes of *nod* remain unknown, but considering the similar effects of *nod* and *Lgn-R/+* on stress-inducible transcriptional programs, we speculate *nod* modifiers might also participate in regulating immunity signaling pathways, which will be a focus of future investigations.

One of the striking results of this study is the contrast at the molecular scale between phenotypic effects in the A619 and B73 inbred backgrounds. From a large-scale view, all of the mutants have comparable effects on growth, transcriptional programs, and metabolism, although they vary considerably in the magnitude of their impact. Fine-scale analysis reveals that the specific genes or metabolites induced or repressed in B73 and A619 are not easily predicted from one background to the next, however (Supplementary Dataset 1). This could be a result of upstream genetic modifiers that reorient signal transduction from NOD and LGN to the nucleus, or could be a consequence of differences in genetic regulatory elements downstream of NOD and LGN. The latter hypothesis is well-supported by the literature: genes involved in specialized metabolism, stress responses, and pathogen defense are rapidly evolving and highly variable within populations, but are typically induced by a core set of common signal transduction pathways (stress signaling “hubs”) (Guo, Major and Howe, 2018; Margalha, Confraria and Baena-González, 2019; Lacchini and Goossens, 2020; Zhang, Zhao and Zhu, 2020). Overall, these findings demonstrate the critical importance of investigating how even severe mutant phenotypes can be differentially expressed in distinct genetic backgrounds, especially in agricultural species with the incredible genetic diversity of maize.

Regardless of genetic background, the *nod;Lgn-R/+* double mutants always exhibited more severe defects than either single mutant, directly demonstrating that *nod* and *Lgn-R* are not epistatic to each other. If NOD and LGN universally acted in a heteromeric complex with each other, or if LGN-catalyzed phosphorylation of NOD were required to mediate signal transduction from LGN, then we would have expected to observe epistasis in the double mutant. Alternatively, if NOD and LGN acted in entirely independent pathways, we might have expected to see partial overlap in effects between each single mutant and the double (purely “additive” effects), but little overlap between the single mutants. Instead, all three genotypes primarily impacted the same biological processes in every phenotype we measured, with coordinate effects on growth, developmental patterning, transcriptional networks, and metabolism. Therefore, we hypothesize that NOD and LGN act on signal transduction pathways that partially converge in maize cells, and propose that NOD and LGN could sometimes (although not always) work in concert at the plasma membrane to sense cues or transduce signals to the cytoplasm.

## METHODS

### Plant growth conditions, genotypes, phenotyping

To generate the double mutant in A619, *Lgn1-R/+* was crossed to A619 and that plant was crossed to homozygous *nod* mutants in A619. *Lgn1-R* plants from this cross were crossed again to *nod* homozygotes in A619 generating family 2431. To generate the double mutant in B73, *Lgn1-R/+* plants were crossed to plants heterozygous for *nod* in B73. *Lgn1-R* plants genotyped to identify the *nod* heterozygotes were crossed again to *nod* heterozygotes generating family 2729.

Family 2431 (A619) was grown in Gill Tract, UC Berkeley for measurements at maturity and grown in the greenhouse in peat pots for RNAseq and metabolomics analysis. Metabolomic analysis was carried out using the second leaf blade at 28 days after sowing. RNAseq was carried out on the same family at 4 weeks after sowing with 1 cm pieces of the shoot apex after removing 3-5 leaves.

Family 2729 (B73) was grown in 2 gal pots in the greenhouse to maturity for photographs and measurements and in peat pots for metabolomics and RNAseq. Metabolomic analysis was carried out using the second leaf blade at 25 days after sowing. RNAseq was carried out on the same plants with 1 cm pieces of the shoot apex after removing 2-4 leaves.

### Coimmunoprecipitation screen

Ten grams of wild type SAM maize tissue were used for each replicate and null alleles were used as a negative control. Membrane proteins were extracted using buffer A [50 mM Tris-HCl, pH 7.5, 150 mM NaCl, 2% IGEPAL-CA-630 (Sigma) and 1× protease inhibitor mix (Roche)]. Protein complex was purified using Dynabeads Epoxy M270 (Thermo Fisher Scientific) with Anti NOD antibody covalently coupled. After four washes with 1XPBS+T, bound target proteins were eluted with a soft elution buffer [50 mM Tris-HCl, pH 7.5, 0.2% SDS, 0.1% Tween 20]. The complex was precipitated with acetone and then separated by SDS-PAGE. Western blot with specific antibodies confirmed the presence of the bait protein. Protein complex was in-gel trypsin digested and analyzed by LC-MS/MS, using a LTQ Orbitrap XL Mass spectrometer (Thermo Fisher Scientific). For protein identification, Uniprot *Zea mays* database in the Mascot package was used. results were exported into Scaffold v4.4.6 (Proteome Software Inc., Portland, OR).

### Yeast two-hybrid screen

The ULTImate Y2H screen was carried out by HYBRIGENICS SERVICES (Paris, France) using a *Zea mays* vegetative SAM, ear/tassel inflorescence cDNA library. The NOD full length cDNA was used as bait and N-terminally fused to the DNA-binding domain in the pB66 vector.

### Bimolecular fluorescence complementation

Full coding sequences of *nod*, *Lgn* and *nod-2* were amplified using specific primers, cloned into the pENTR D vector (Invitrogen), and transferred to BiFC vectors pB7WGYN2 or pB7WGYC2 by LR recombination. *Agrobacterium tumefaciens* strain GV3101 was transformed with BiFC constructs. Agroinfiltration into *Nicotiana benthamiana* leaves was performed as described previously (Bolduc and Hake, 2009). Twenty four hours after agroinfiltration, leaves were observed under LSM710 confocal microscopy (Zeiss) with 470-nm excitation and 535-nm emission filters.

### *In vitro* kinase experiments

To produce recombinant NOD, LGN, and Lgn-R protein, cDNAs corresponding to the full length with no transmembrane domain were amplified and cloned into pDEST15 (Invitrogen). *E. coli* BL21 was transformed with constructs, and GST tagged proteins were induced in 500 mL of media for 3 hours with 1 mM isopropylthio-B-galactoside (IPTG) when OD_600_ was 0.5. Bacterial pellet was collected by centrifugation and lysed using B-PER™ Protein Extraction Reagent (Thermo Fisher Scientific). Soluble protein was purified with 200 μl of Glutathione agarose beads slurry (Sigma), gently shaking for 1 hour at 4°C. Beads were washed five times with 1X PBS + Tween 20. Purified proteins were eluted with 10mM reduced glutathione buffer. For kinase assays, 10μg of protein in 10 μl was mixed with 2 μl 10X kinase buffer (125 mM Tris, pH 7.5, 25 mM MgCl2, 1 mM EDTA), 1 μL [γ- P] ATP (3000 Ci/mM) and H2O, and incubated for 30 minutes at 25°C. SDS- PAGE loading buffer (250 mM Tris-HCl pH 6.8, 2% SDS, 30% glycerol, 0.1 M DTT, 0.02% Bromophenol Blue) was added to each sample to stop the reaction. Samples were boiled and separated on 12% polyacrylamide gels. Autoradiography was carried out for detection. For fluorescent detection, cold ATP was used in the kinase assays at 200 μM and detection was using ProQ diamond stain (Invitrogen).

### Protein mass spectrometry

The mass spectrometry instrument used to analyze the samples was a Xevo G2 QTof coupled to a nanoAcquity UPLC system (Waters, Milford, MA). Samples were loaded onto a C18 Waters Trizaic nanotile of 85 um × 100 mm; 1.7 μm (Waters, Milford, MA). RAW files were processed using Protein Lynx Global Server (PLGS) version 2.5.3 (Waters, Milford, MA). For viewing, PLGS search results were exported into Scaffold v4.4.6 (Proteome Software Inc., Portland, OR).

### RNA-Seq Analysis

RNA-Seq was performed as previously described (Scarpin, Leiboff and Brunkard, 2020; Busche *et al*., 2021). Briefly, sequenced reads were aligned to the maize genome (B73 Refgen V3) with HISAT2 (Kim, Langmead and Salzberg, 2015) and counted with HTSeq (Anders *et al*., 2015). Differential transcript abundance was determined with DESeq2 (Love, Anders and Huber, 2014). Functional analysis of differentially-expressed genes was performed with MapMan gene annotation (Thimm *et al*., 2004; Usadel *et al*., 2009; Lohse *et al*., 2014).

### Untargeted metabolomics

Ground and frozen tissue (50 mg) in 1.5-mL fast prep tubes was carefully thawed to 4°C then transferred to ice. An internal standard mix of 12 μL containing caffeine, D6-Abscisic acid, D5-Jasmonic acid, D5-Cinnamic acid, D5-Indole-3-acetic acid, 13C-alpha linolenic acid, and nicotine at 8.33 μg/mL was added and plant metabolite extractions were performed as described (Christensen *et al*., 2021). Ultra-high- performance liquid chromatography-high-resolution mass spectrometry (UHPLC-HRMS) was carried out on both a Q-Exactive and Fusion mass spectrometer coupled to a Vanquish LC System (Thermo Fisher Scientific, Waltham, MA, USA) by reverse phase gradient elution using an ACE Excel 2 C18-PFP column (2.1 μm, 100 mm) in full scan, ddMS2, and ddMSn in positive (injection volume 2 μL) and negative (injection volume 4 μL) ion modes, and the data were acquired, processed, normalized, filtered and metabolites identified using MZmine 2 (Pluskal *et al*., 2010) and Metaboanalyst 4.0 (Chong *et al*., 2018) software as described (Christensen *et al*., 2021).

The benzoxazinoids MBOA (NIST 2000; Sigma aldrich 2015), HDMBOA (Glauser *et al*., 2011; Tsugawa *et al*., 2019) and HDMBOA-Glu (Glauser *et al*., 2011; Marti *et al*., 2013), were detected in positive mode at eV 20, 35, 50, 100 (MBOA), eV 20, 35, 50 (HDMBOA), and eV 20, 50, 100 (HDMBOA-Glu). Comparative spectral referencing for these compounds was performed using MS2 diagnostic fragmentation peaks (Glauser *et al*., 2011; Marti *et al*., 2013; Tsugawa *et al*., 2019), RT, and accurate mass data available on the MassBank, NIST, mzCloud, and GNPS databases. Mass Frontier 8.0 (Thermo Fisher Scientific, Waltham, MA, USA) was used to accurately annotate and assign fragmentation structures of the MSn spectral data, and the Mass Frontier fragmentation library and Fragment Ion Search function (FISh) was utilized to estimate and assign metabolite secondary fragmentation structures. The mzLogic function was also used to further confirm fragment annotation though matching individual MSn substituent groups to existing substituent fragmentation spectra via the mzCloud database. Documentation on MSn fragmentation annotation can be found in supplementary tables (Supplementary Dataset 2).

## Supplemental Materials

**Supplementary Dataset 1**. Transcriptomes and metabolomes of single and double mutants in B73 and A619 backgrounds.

**Supplementary Dataset 2**. MSn fragmentation annotation.

## Acknowledgments and Funding

We thank Samuel Leiboff and Alyssa A. Anderson for assistance with experiments and analyses. This work was funded by the National Institutes of Health OD DP5-023072 to J.O.B., the National Science Foundation ECA-PGR 1733606 to S.H., and the UC-Mexus CONACYT collaborative grant CN-19-149 postdoctoral fellowship to M.J.A.-J. and S.H.

## Author Contributions

M.J.A.-J., M.B., C.L., S.H., and J.O.B. designed the research; M.J.A.-J., M.B., C.L., J.W., S.A.C., C.T.H., S.H., and J.O.B. performed research; J.W., S.A.C., and C.T.H. contributed analytic tools for quantifying metabolite levels; M.J.A.-J., M.B., C.L., J.W., S.A.C., C.T.H., S.H., and J.O.B. analyzed data; and M.J.A.-J., M.B., C.L., S.H., and J.O.B. wrote the manuscript.

